# Multi-Model and Network Inference Based on Ensemble Estimates: Avoiding the Madness of Crowds

**DOI:** 10.1101/858308

**Authors:** Michael P.H. Stumpf

## Abstract

Recent progress in theoretical systems biology, applied mathematics and computational statistics allows us to compare quantitatively the performance of different candidate models at describing a particular biological system. Model selection has been applied with great success to problems where a small number — typically less than 10 — of models are compared, but recently studies have started to consider thousands and even millions of candidate models. Often, however, we are left with sets of models that are compatible with the data, and then we can use ensembles of models to make predictions. These ensembles can have very desirable characteristics, but as I show here are not guaranteed to improve on individual estimators or predictors. I will show in the cases of model selection and network inference when we can trust ensembles, and when we should be cautious. The analyses suggests that the careful construction of an ensemble – choosing good predictors – is of paramount importance, more than had perhaps been realised before: merely adding different methods does not suffice. The success of ensemble network inference methods is also shown to rest on their ability to suppress false-positive results. A Jupyter notebook which allows carrying out an assessment of ensemble estimators is provided.

## 1 Introduction

In physics simple and elegant symmetry relationships have often led the way to theoretical models (Thorne & Blandford, 2017). Most importantly Emmy Noether’s theorem has been pivotal in establishing the correspondence between continuous symmetries and conservation laws in physics (Neuen-schwander, 2017); it has been pivotal in the derivation of physical *laws of nature*. Biology has not been able to draw on such fundamental principles (May, 2004), to a large degree because most processes are intrinsically dissipative (meaning energy is produced andconsumed) and hence the conditions where Noether’s theorem holds simply do not apply. Instead biological models have often had a heuristic element, or described the rates of change in (often macroscopic) observables (e.g. species of plants or animals, or molecular species).

Writing down the set of equations is an important starting point in modelling as it forces us to express our assumptions in a precise form. Which form to choose is, however, not unambiguously obvious. Instead we often rely on data to decide between the different options. *Statistical model selection* (Burnham & Anderson, 2013) provides the tools to make such decisions, balancing the ability of a model to fit the data with the model’s complexity. As larger and larger models, even models for whole cells (Karr *et al*., 2012, 2015; Lang & Stelling, 2016; Babtie & Stumpf, 2017), are being considered model selection problems will presumably become the norm, especially when models are constructed exhaustively or automatically (Ma *et al*., 2009; Barnes *et al*., 2011; Szederkényi *et al*., 2011; Sunnåker *et al*., 2014; Babtie *et al*., 2014; Leon *et al*., 2016; Gerardin *et al*., 2019).

In some, probably many, situations model selection will not be able to decide on a single best model. Instead many models may have comparable support. In such a situation we may then base analysis or predictions on the ensemble of models that are supported by the data. Each model’s contribution to the prediction etc. is weighted by the relative support it has received. Such estimates or predictions based on ensembles have been referred to as exploiting the “*wisdom of crowds*” (Marbach *et al*., 2012). This refers to the notion that groups of individuals/models are more likely to the be better than those based on a single individual/model. This concept, however, also references a much earlier work, Charles Mackay’s “*Extraordinary Popular Delusions and the Madness of Crowds*” (Mackay, 1841), a 19th century account of how popular opinion can support quite extraordinary and plainly wrong opinions and concepts.

Ensemble estimators have a surprisingly long history, outlined in Laan *et al*. (2017), and aspects such as bagging, boosting, and stacking (Murphy, 2012) are firmly established in the statistical learning literature; see for example Kuncheva (2004) for a comprehensive treatment. There has been interest in evolutionary biology (Strimmer & Rambaut, 2002); and following Marbach *et al*. (2012), there been further developments in systems and computational biology e.g. (Saez-Rodriguez *et al*., 2016; Thorne & Stumpf, 2012). But in the context of network inference, combining different network reconstruction methods has sometimes been viewed as necessarily optimal (Le Novère, 2015). Below we show that this is not automatically the case. In turn, I will show that model averaging and ensemble estimation are susceptible to poorly defined sets of candidate models; and that the behaviour of ensemble approaches to network reconstruction depends strongly on the composition of the ensemble. For very good ensembles, the advantage comes mainly from reducing the number of false-positive edges. Both problems share in common a dependence on the quality of the ensemble, and we map out and quantify these influences; we also provide self-contained Julia code for further *in silico* experimentation and analysis of ensemble prediction methods.

## 2 Model Selection and Multi-Model Inference

We assume that we have a *universe* of models, ℳ,

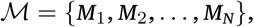

that are potential mechanisms, by which some data, 𝒟, have been generated. For simplicity, we consider a finite number of models, *N*. Furthermore, for each model, *M*_*i*_, we assume that we have a data generating function, *f*_*i*_(*θ*_*i*_), parameterised by a parameter vector *θ*_*i*_ which is chosen from some suitable continuous parameter space,

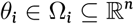

Coping with the size of the parameter space (Cox, 2006) is one of the essential challenges of parameter estimation and model selection.

We start from a Bayesian framework (Robert, 2007), where we seek to determine the *posterior distribution* over parameters,

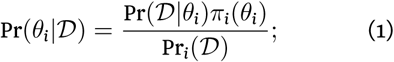

where Pr(𝒟 | *θ*_*i*_) is the *likelihood*; *π*_*i*_(*θ*_*i*_) the *prior* over the parameters for model *M*_*i*_, and Pr_*i*_(𝒟) = ∫Pr(𝒟 | *θ*)*π*_*i*_(*θ*)*dθ* (here we make the dependence on the choice of model explicit through an index) is known as the *evidence*.

In the Bayesian framework model selection is a (relatively) straightforward extension, and the *model posterior* is given by

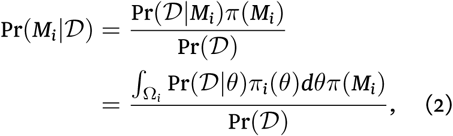

where analogously to Eqn. (1), we have the *model posterior* Pr(*M*_*i*_|𝒟), and *model prior π*(*M*_*i*_). The denominator terms in Eqns. (1) and (2) are notoriously hard to evaluate for all but the simplest cases, and a large amount of ingenuity and work has been invested into computational schemes(Green *et al*., 2015), including *Markov Chain Monte Carlo, Sequential Monte Carlo*, and related approaches. Often even the likelihood is prohibitively expensive to evaluate and so-called *approximate Bayesian computation* (ABC) schemes have been devised to make Bayesian statistical inference possible (Toni & Stumpf, 2010).

Alternatives to the Bayesian framework, such likelihood-based inference(Cox, 2006) and optimisation of cost-functions (Raue *et al*., 2014), result in *point estimates* for the parameters, e.g. the value of *θ*′ that maximises the probability of observing the data,

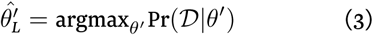

Similarly, we can determine the *maximum aposteriori estimate* by finding the mode of the posterior distribution (Robert, 2007; MacKay, 2003),

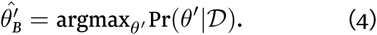

Compared to analysis of the posterior distribution, such local estimates loose a lot of relevant information, but some characteristics can still be recovered by considering the local curvature of the likelihood, i.e. the *Fisher Information*, or cost-function surface around the (local) extremum identified in this manner (Gutenkunst *et al*., 2007; Erguler & Stumpf, 2011).

Model selection frameworks that are based on likelihood inference rely on criteria to find the optimal model among a set of candidate models. The *Akaike Information criterion* (AIC) (Akaike, 1974; Burnham & Anderson, 2013) for model *M*_*i*_ is given by

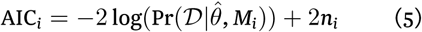

with 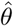 given by Eqn. (3), and *n*_*i*_ the number of parameters of model *M*_*i*_. The AIC is probably the most widely used model selection criterion, despite the fact that it is biased in favour of overly complicated models as the amount of available data increases. The *Bayesian information criterion* does not suffer in the same way; it is defined as

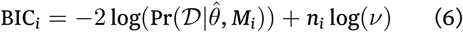

with *ν* representing the size of the data or number of samples. Several other information criteria exist (discussed e.g. in (Burnham & Anderson, 2013; Kirk *et al*., 2013)), but they all share in common the purpose of balancing model complexity with model fit. The BIC can be derived as an approximation to the full Bayesian model posterior probability, which achieves this balance implicitly.

If model selection cannot pick out a clear winner, then either (i) further analysis should be used to design better, more informative experiments that can discriminate among these models(Liepe *et al*., 2013; Busetto *et al*., 2013; Silk *et al*., 2014); or (ii) these models should be considered as an ensemble (Burnham & Anderson, 2013; Thorne & Stumpf, 2012). The former has definite attractions as it will lead to an increase in our understanding if we can discard some of the models/mechanistic hypotheses.

The latter approach, basing analysis and especially predictions on an ensemble of models has become a popular concept in machine learning. Most notably, in the context of biological network inference the “*Wisdom of Crowds*” concept (Marbach *et al*., 2012) has been important in popularising inference based on several models. Here we are considering model averaging where contributions from different models may be weighted by their relative explanatory performance. In the Bayesian framework (Hoeting *et al*., 1999) we can use the posterior probability directly. In the context of an information criterion *I*_*i*_ formodel *i* we define (Burnham & Anderson, 2013)

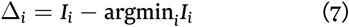

and then determine the model weight as

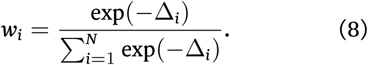

The model weights (e.g. the Akaike weight if *I*_*i*_ is the AIC) provide the relative probability of model *M*_*i*_ to be the correct model conditional on the data 𝒟 and the set of models, ℳ, considered. Model averaging in this framework can serve as a *variance reduction technique* (Burnham & Anderson, 2013; Murphy, 2012).

## 3 Statistical Physics of Model Selection and Ensemble Estimation

In order to simplify the discussion we define a relationship between the model probability (always implicitly understood to be either a posterior or relative model probability), *p*_*i*_, and a *cost* or *energy, ϵ*_*i*_, as (Babtie & Stumpf, 2017)

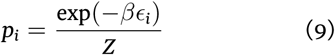

with the normalisation constant *Z* (the *partition function*) given by

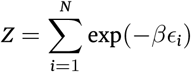

With this in hand we can straightforwardly consider different model selection/averaging frameworks in a similar manner.

In general the true data-generating model (a natural system) will not be represented perfectly by any of the models in ℳ; we denote this true model by ℵ ∉ ℳ. But if we are interested in finding out whether has a certain characteristic *ϕ* we would obtain this from the appropriate ensemble average

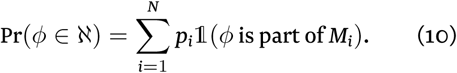

where 𝟙() is the conventional indicator function, i.e.

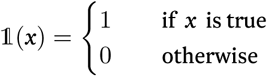

Eqn. (10) is based on three, potentially strong assumption.

1. The model universe, ℳ, is “complete” in the sense that we expect ℳ to contain any model, *M*_*k*_, which we might expect to be a reasonable description of ℵ (always remembering that ℵ ∉ ℳ.
2. It is decidable if *ϕ* is part of *M*_*i*_, ∀*M*_*i*_ ∈ M.
3. *ϕ*, or any *ϕ* of interest has played no part in the construction of the model probabilities, *p*_*i*_, Eqn. (9).

The first assumptions is arguably the strongest assumption. In principle we can update ℳ → ℳ′ by adding additional models in light of data; specifying new priors, *π*′ for the models 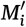 ∈ ℳ′, will require some care. One important condition that ℳ (and ℳ′) must fulfil is that models must not appear more than once in the set. This is important to keep in mind as we are increasingly relying on automated generation or exhaustive enumeration of models.

With fixed 𝒟 and ℳ Eqn. (10) is, however, a good approach for ensemble-based prediction and estimation. It also encompasses Bayesian model averaging(Hoeting *et al*., 1999) and multi-model inference based on information criteria (Burnham & Anderson, 2013).

### The effective model space

We first analyse a simple case where all models in our universe have associated costs drawn from a suitably well behaved probability distribution, *q*(*η*) (Babtie & Stumpf, 2017), with positive support and associated density, *f*_*q*_(*ϵ*), over the model energies, *ϵ*_*i*_, such that

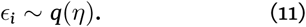

Because *ϵ*_*i*_ is a random variable, the relative weight, *ω*_*i*_ = exp(− *ϵ*_*i*_), will also be a random variable, and we can obtain the probability density function, *f*(*ω*), via change of variables as

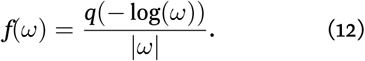

For different choices of *q*(*ϵ*) we can now investigate the distribution over model weights. For example if *ϵ*_*i*_ ∼ Gamma(*α, θ*) (where *α* and *θ* denote the shape and scale parameters, respectively) then

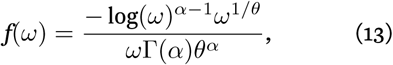

with Γ(.) the Gamma function. The Gamma distribution is a flexible and generic distribution and is chosen for its generality rather than any particular property and our discussion here does not depend on its specifics. Some representative distributions over *ϵ* and the corresponding distributions over *ω* are shown in Fig. 1, panels (A) and (B).

**Figure 1:**
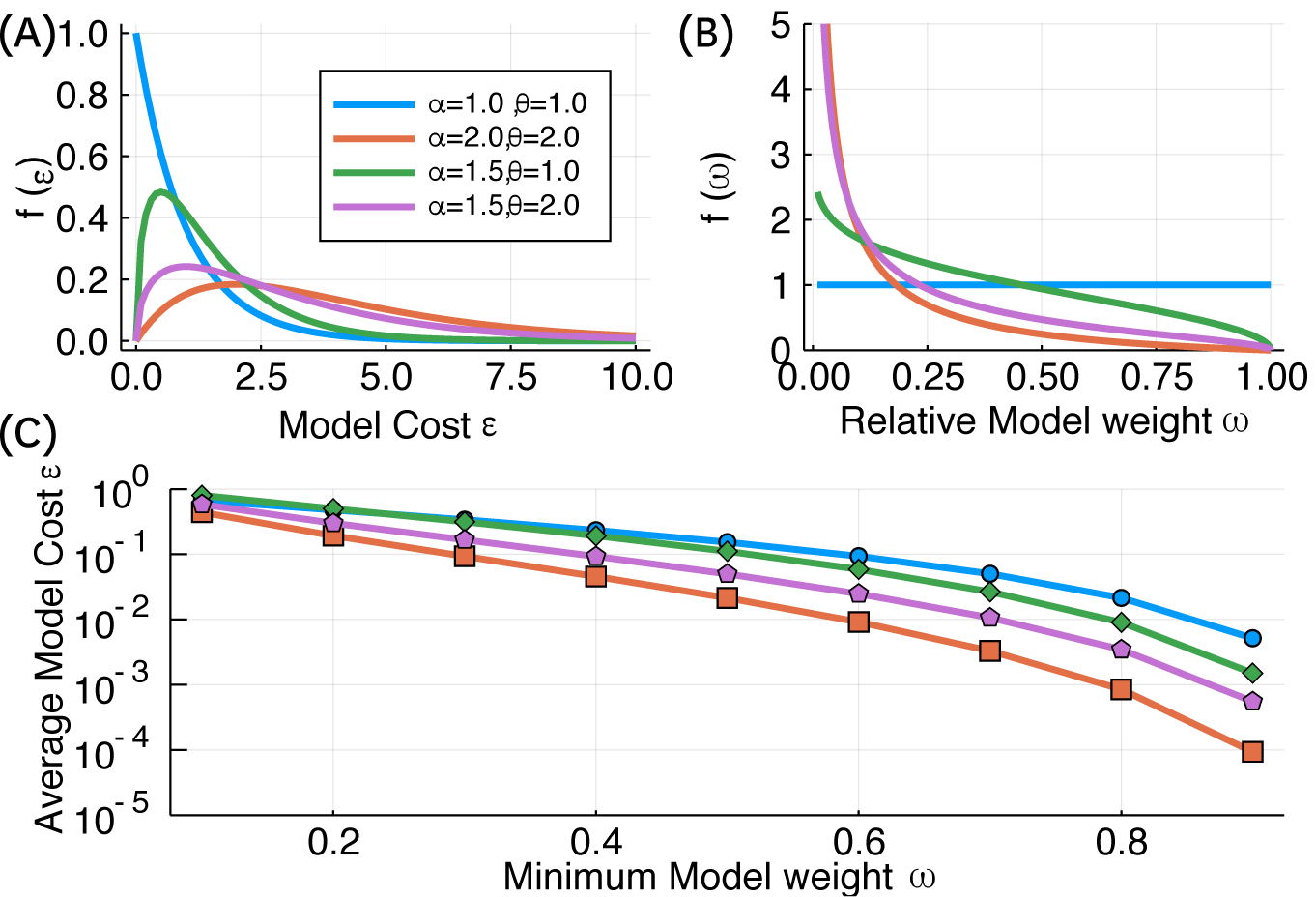
Different distributions over the model costs, ϵ_i_ (top left) and the resulting distributions over the relative model weights (top right). Models seen as good (low cost) correspond to those with higher weights, ω. Note that for α = θ = 1 the Gamma distribution reduces to an Exponential distribution, and therefore the distribution over model weights becomes uniform. The bottom panel shows the average cost, ϵ, if models above a given minimum relative weight are included.

The distribution over model costs, *ϵ*, affects the distributions over model weights, *ω*. This is important to realise when deciding on how to triage model sets for further analysis and prediction (Cade, 2015): generally, inference based on all models weighted by *ω*_*i*_ is neither practical nor desirable, as many models with low weight will mask the information available in the good models. If, for example, we only include models with *ω* ∈ [0.9, 1.0] then the average model cost

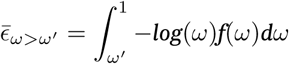

(because *ϵ* = − log(*ω*)) for these sets will be 0.005 (blue), 0.0015 (green), 6×10^*−*4^ (purple), and 9×10^*−*5^ (orange).

Fig. 1(C) shows that the (unknown) distribution over costs can affect multi-model inference quite profoundly. But for model universes that are enriched for good models (i.e. many models *M*_*i*_ with low values of *ϵ*_*i*_) selecting a subset of models based on even a fairly conservative threshold for the model weights *ω*_*i*_, can result in a sufficiently large model sets for further prediction.

### A simple test case for multi-model inference

Here we study a very simplistic scenario in which we have three types of models, borrowing and adapting Box’s (Box, 1976) terminology:

### Useful Models

which capture important aspects of ℵ and which have an associated cost *ϵ*_1_.

### Useless Models

which are qualitatively and quantitative poor descriptions of reality and have an associated cost *ϵ*_2_ ≫ *ϵ*_1_.

### Nuisance Models

which are qualitatively different from reality, but which can quantitatively capture aspects of ℵ by chance; their costs are random variables, *η* ∼ [0, *τ*].

Nuisance models are here assumed to be models where the quantitative agreement with data is unrelated to the data generating mechanism ℵ. Purely machine-learning based models are one way in which we can realise such nuisance models (Camacho *et al*., 2018); for small datasets 𝒟, poorly designed experiments, or simply lack of prior knowledge, there are many ways in which model fit may only be a poor reflection of reality, and this can also give rise to nuisance models (Baker *et al*., 2018).

For concreteness let these three different model classes have sizes *ν*_1_, *ν*_2_ and *ν*_3_ = *N* − *ν*_1_ − *ν*_2_, and assume that 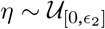 i.e. nuisance models are at worst as bad as useless models. Then the fraction of nuisance models that have lower associated costs than the *useful models* is given by

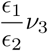

The relative influence of nuisance models can be studied by contrasting three features, *ϕ*_1_, *ϕ*_2_, and *ϕ*_3_, with the following properties

*ϕ*_1_ : equally represented with frequency *ξ* among models of all classes.

*ϕ*_2_ : only represented among useful models.

*ϕ*_3_ : only represented among “nuisance” models.

With Eqn. (10) we can obtain Pr(*ϕ*_*i*_ ∈ ℵ) for any property with frequencies *ξ*_*i*_ in the classes *i* = 1, 2, 3,

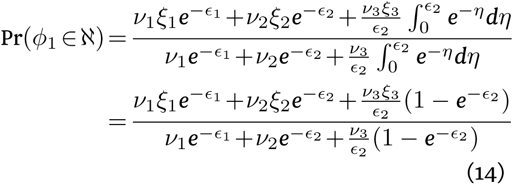

First, for *ϕ*_1_ we trivially obtain

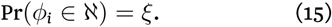

For the more interesting probability for *ϕ*_2_ under the model averaging framework we obtain,

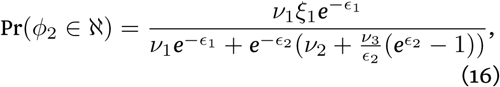

and finally, for a characteristic shared by and confined to the set of nuisance models, we obtain,

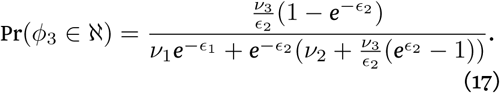

From Eqns. (16) and (17) we see that multimodel averaging is prone to propagate wrong results if nuisance models are frequent and some receive good quantitative support (i.e. low model costs, *ϵ*). Equally, worrying, the same scenario can make it hard for true aspects of ℵ to receive sufficient support through Eqn. (16) if there are many nuisance and useless models included in the data.

To illustrate this further we can consider the case where *ξ*_2_ = 1 and *ϕ*_3_ = 1 (meaning every useful model exhibits characteristic *ϕ*_2_, and every nuisance model characteristic *ϕ*_3_) and ask when is Pr(*ϕ*_3_ ∈ ℵ) *>* Pr(*ϕ*_2_ ∈ ℵ)? We obtain

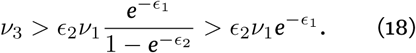

Thus if useful models are sufficiently rare in the model set (say *ν*_1_ *<* 0.1) the nuisance models’ characteristics will have high weight in the ensemble average, see also Figure 2. Neither of the parameters, *ν*_1_, *ν*_2_, *ν*_3_, *ϵ*_1_, *ϵ*_2_ are, of course, known, and we cannot know which class a model belongs to *a priori*.

**Figure 2:**
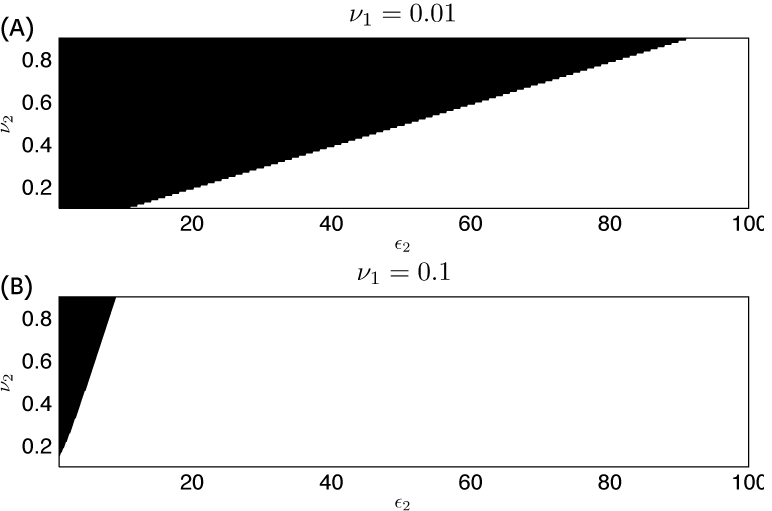
For ϵ_1_ = 0.1 we indicate the areas in the ϵ_2_, ν_2_ space, where the contribution to ensemble estimators coming from the nuisance models exceeds that coming from the useful models (black). On the top the frequency of useful models is ν_1_ = 0.01; in the bottom panel we set ν_1_ = 0.1.

Thus model averaging is not a panacea and requires careful consideration of which models are included in the set.

## 4 Ensemble Estimation and Network Inference

Network inference can also benefit from ensemble methods (Marbach *et al*., 2012; Meyer *et al*., 2014; Saez-Rodriguez *et al*., 2016) but here, too, potential pitfalls can arise. We are considering directed networks (Kolaczyk, 2009; Koller & Friedman, 2009), with *V* nodes, and *L* edges; the *adjacency matrix, A*, is a convenient way to specify such networks, if we indicate the presences and absences of edges, by one and zero, respectively, in its entries,

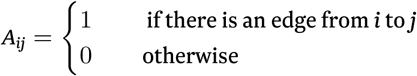

In network inference we seek to determine whether the data 𝒟 suggests statistical relationships among pairs of nodes *v*_*i*_, *v*_*j*_ that may indicate a functional relationship, represented by an edge connecting the nodes. We consider *k* different algorithms, *O*_*κ*_*κ* = 1, …, *k*, which predict the presence, 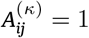 or absence 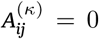, of a link from *i* to *j*. If we have for the false positive and false-negative probabilities for method *κ*,

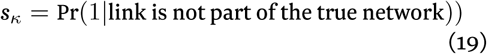

And

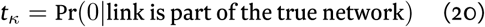

we can assess how beneficial the ensemble estimators are for the quality of the inferred networks.

### Ensembles of Identical Estimators

The simplest case, which is already instructive as a baseline, is where all methods have identical performance, *s*_*κ*_ = *s* and *t*_*κ*_ = *t*, ∀*κ* = 1, …, *k*. If the performance of the inference methods is statistically independent, then the number of agreeing inference methods is a binomial random variable. If we set a threshold on the minimum number of concordances among inference methods, *k*_0_, we then have for the overall probability of scoring a true edge from the ensemble,

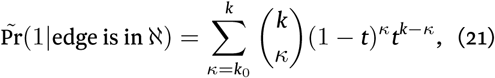

while the probability of a false negative is

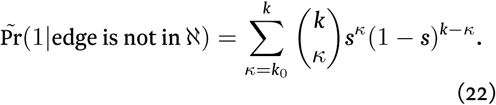

To illustrate the outcome of such a simple estimation procedure we assume a network loosely based on expected *Homo sapiens* network sizes(Stumpf *et al*., 2008) (22, 000 nodes and 750, 000 interactions). In figure 3 we consider 10 ensembles and base the ensemble estimator on the minimum number, *k*_0_, of methods that have to predict an edge. If *k*_0_ is too small then there will be too many positives as is the case here; a *majority-vote* vote, i.e. here *k*_0_ = 6, would do an acceptable job in terms of precision, recall and F1 statistic (Murphy, 2012), but this does depend on the false positive ratio in particular (as biological networks are sparse), as well as the size of the ensemble of inference algorithms, 𝒪 = {*O*_1_, *O*_2_, …, *O*_*k*_}.

**Figure 3:**
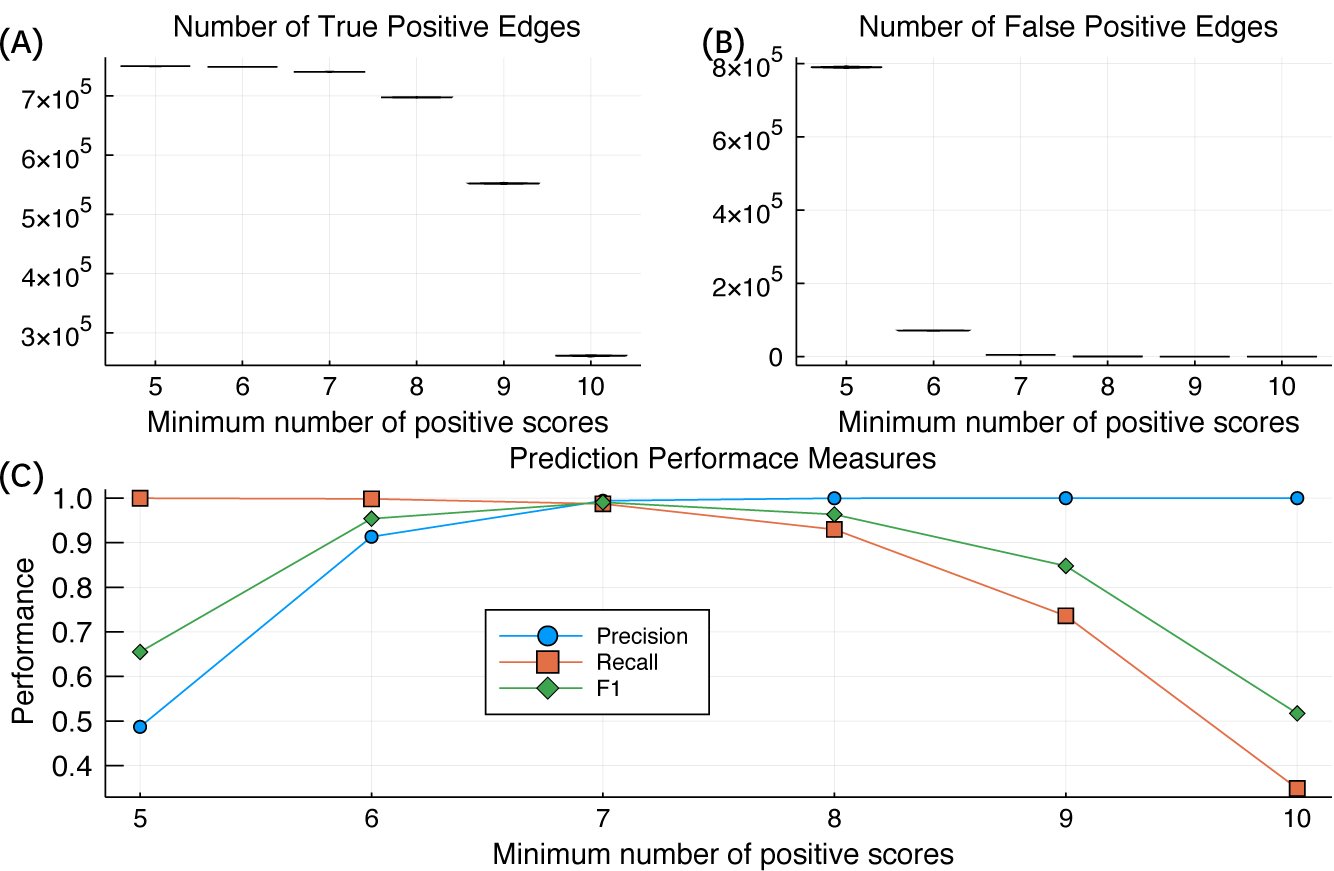
Illustration of the performance of ensembles of network inference methods with with false positive and false negative probabilities s = t = 0.1. Real biological networks are sparse unless false-positives are controlled in the ensemble(Stumpf & Wiuf, 2010; Penfold & Wild, 2011); once false-positives are over-controlled (by demanding a larger number of methods to score an edge), the recall deteriorates.This is reflected in the top panels (the number of true and false positives (we generated 1000 random inferred networks but because of the large network size and the binomial nature of the process the distributions around the mean are tight. The lower panel shows the precision, recall and F1 statistics (Murphy, 2012; Mc Mahon et al., 2014) (see also Appendix A) as a function of the minimum number of methods that need to positively score an edge (again confidence intervals are very tight to be effectively hidden by the symbols.

### Ensemble Estimators can be Worse than Individual Estimators

We are interested in ensemble estimators because we know that individual estimators are far from perfect.

But ensembles are not guaranteed to always improve on individual estimators. We compare an ensemble of equally well performing estimators with a single estimator. The ensemble false negative probability, *T*, is given by

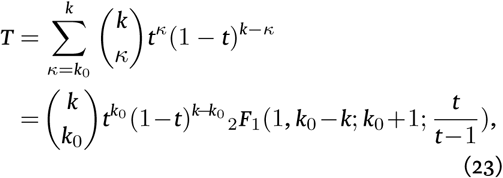

where _2_*F*_1_ is the Hypergeometric function (Arfken *et al*., 2013)(see Appendix Bfor an approximation for small arguments). From this we can determine when the ensemble false negative rate, *T*, will be greater than *t*,

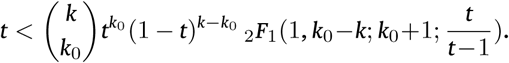

Equally, we obtain for the ensemble false positive rate, *S*,

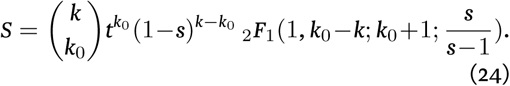

For low thresholds, *k*_0_, the ensemble error rate can be greater than that of the individual estimator; this is because the stringency of the ensemble prediction is then reduced, as the cumulative probability of a sufficiently small number of estimatorsto “accident ally” agree is greater than the error rate of the individual estimator. We show this for two false negative rates, *t* = 0.1, and *t* = 0.05, in figure 4 (A) and (B).

**Figure 4:**
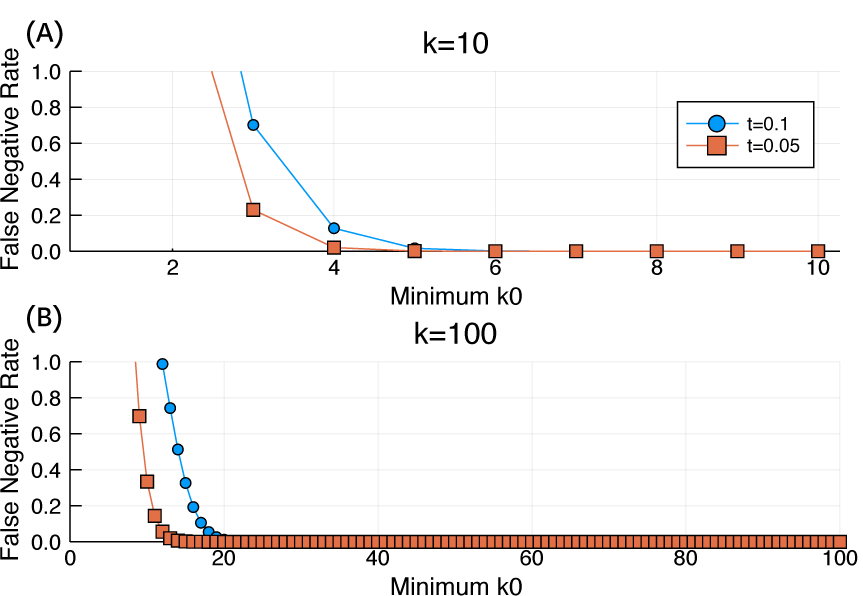
Illustrative cases where the error rate of an ensemble estimator exceeds the error rate of an individual estimator for k = 10 and k = 100. Reassuringly, this only happens for small values of the threshold k_0_.

### 4.1 Heterogeneous Ensembles of Network Inference Methods

We next focus on the case of a small set of predictors, *k* = 10, and two classes of methods: a set of good methods with error rates *t*_1_ and *s*_1_; and a set of poor methods with error rates *t*_2_ *> t*_1_ and *s*_2_ *> s*_1_. In figure 5 we show, again for a case modelled on a likely human gene regulation network, the likely true and false positive predictions arising from ensembles with different numbers of good vs poor edge prediction methods. The basic lesson is that the good methods have to outnumber the bad methods, otherwise especially the precision will suffer. Here we have chosen a simple majority-vote criterion, and to bring precision and F1 statistic to a satisfying level (say in excess of 0.7) requires essentially purging the ensemble of the weakly performing methods (i.e. *k*_1_ ≳ 8). This only points to the extent to which poor methods can compromise the performance of ensemble estimators (and the accompanying `Jupyter` notebook can be used to explore this further).

**Figure 5:**
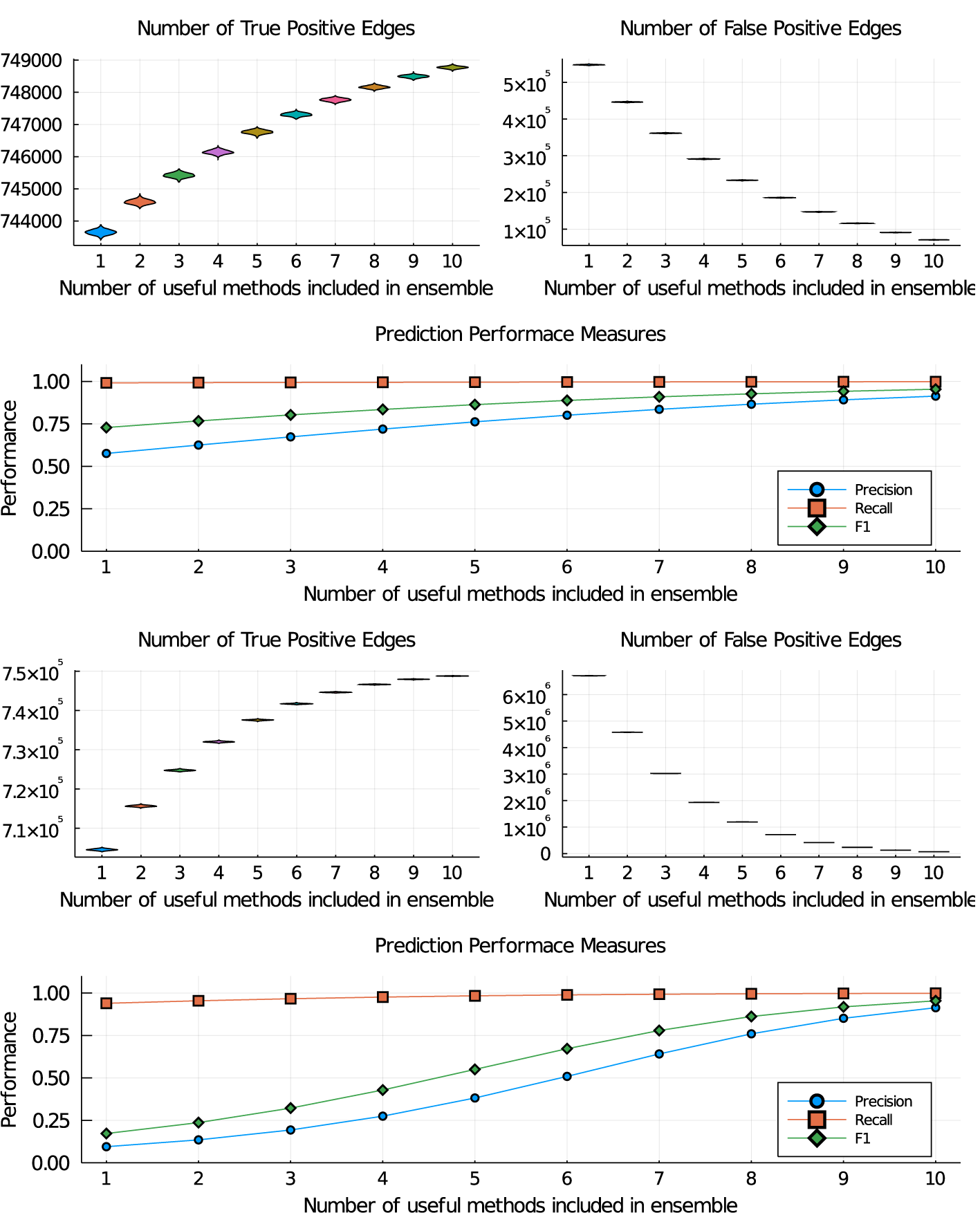
Illustration of the performance of ensembles of network inference methods with with false positive and false negative probabilities s = t = 0.1 for the good predictors and with s = t = 0.15 (panels (A)-(C)) and s = t = 0.25 (panels (D)-(F)). Real biological networks are sparse and unless false-positives are controlled in the ensemble; once false-positives are over-controlled (by demanding a larger number of methods to score an edge), the recall deteriorates.This is reflected in the top panels((A), (B) (D) and (E), showing the numbers of true and false positives(we generated 1,000 random inferred networks). The lower panels((C) and(F)) showthe precision, recall and F1 statistics (Murphy, 2012) (see also Appendix A) as a function of the minimum number of methods that need to positively score an edge. Increasing the false-positive and false-negative error rates from 0.1 to 0.25 for the poor estimators results in marked deterioration of the ensembles. And even small number of bad predictor can profoundly affect the ensemble performance.

**Figure 6:**
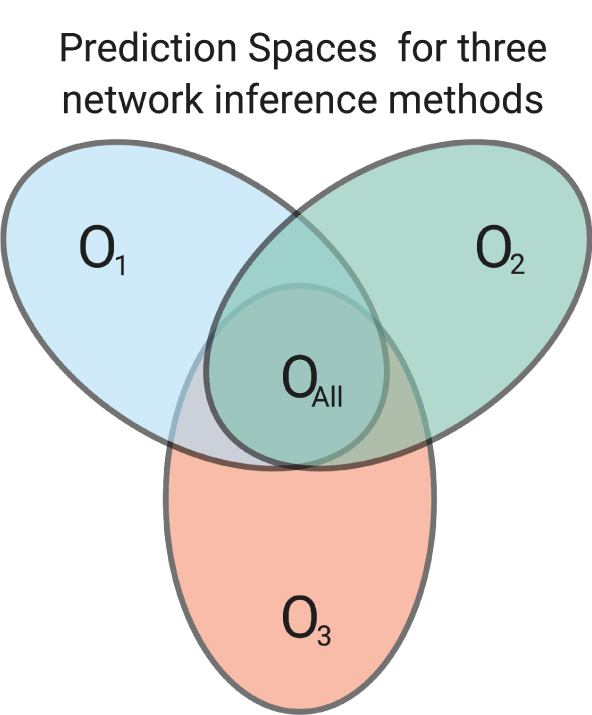
Illustration of different predictors which capture different aspects of the data. If predictors, O_1_, O_2_ and O_3_ have particular performance preferences for certain types of interactions then combining them may improve the ensemble estimation. But this depends crucially on the false-positive rate. In an ideal scenario we would also be able to exploit the individual strengths of the predictors to reduce the ensemble false negative rate; the mathematical formalisms exist(Kuncheva, 2004; Madirolas & de Polavieja, 2015; Díez-Pastor et al., 2015; Laan et al., 2017), but we need to be able to quantify the individual predictors’ behaviours better.

In sparse networks especially, poor estimators will result in inflation of false positive results, and lead to overall poor performance: in a directed network there are *N* × (*N* − 1) interactions possible, but this is still vastly greater than the number of existing edges. For example, in the case of the human network we would have on the order of 4.8 × 10^8^ potential interactions of which only some 750,000 are expected to exist. So even for *s*_2_ = 0.5 there would be about 470,000 cases where ten such methods would agree and score an edge (and 1.8 × 10^8^ if a majority vote rule were applied).

## 5 Discussion

We have shown that ensemble estimators are not as robust as has sometimes been claimed (Le Novère, 2015), or incorrectly surmised from the success of community average results in the DREAM competition (Marbach *et al*., 2012); there, of course, it had already been shown that certain, carefully selected, subsets of estimators give better results than others (Saez-Rodriguez *et al*., 2016).

For the analysis of multi-model inference from mechanistic models we can distill two points: (i) ensembles of mechanistic models that are reasonably defined (Aijo & Bonneau, 2016) (i.e. their construction incorporates any available mechanistic insights; duplicate models are avoided; the model is predictive and can be used to generate data that can be compared with real observations/data) can be combined with the aid of model selection criteria or Bayesian posterior model probabilities with relative ease and safety; (ii) the inclusion of “nuisance models” can hamper ensemble approaches if they come to predominate the model universe ℳ. Such situations could become more likely as model spaces are explored exhaustively(Scholes *et al*., 2019) or automatically (Sunnåker *et al*., 2014). Because of the formalism connecting different model selection criteria, Eqn. (9), these are general results, and do not depend on the particular model averaging procedure chosen (as also clear from the analysis in Laan *et al*. (2017)). So in essence the construction of the models in ℳ (May, 2004; Kirk *et al*., 2015a) will determine the robustness of model averaging or ensemble approaches to prediction and analysis. Little is to be gained by increasing the size of ℳ beyond the (already quite large) set of reasonable models.

In the context of network inference the situation is similar. We find that the poor performance of some methods can drag down the performance of an ensemble estimator or network predictor. So like in the construction of the model universe ℳ before, the make-up of the ensemble of network inference methods, *O*_*κ*_*κ* = 1, …, *k*, does matter considerably (as was also found in the empirical study of Marbach *et al*. (2012)). Majority vote will typically be a sound criterion for a set of reasonable estimators, though not necessarily optimal from the perspective of precision and F1 measures. This is because biological networks are sparse and false-positives will predominate the inferred networks unless they are carefully controlled. For a set of statistically similar powerful inference methods, a conservative criterion for scoring edges will improve on the overall performance of the individual estimators, however.

The problem of network inference has long been known to be challenging. One reason for this (in addition to the large-scale testing problem) is that we do not have a fair way to score and compare the performance of different network inference approaches. The most promising existing approaches are typically computationally expensive and rigorous *in silico* assessment of performance as well as the factors influencing performance is often seen as computationally prohibitively expensive. There is also a danger of biasing simulated data in favour of a given method and the DREAM competition has aimed — with some success we feel — to avoid this, and other approaches have followed suite. Clearly more effort in this domain is needed and computational cost should not preclude rigorous assessment (Chan *et al*., 2017); this situation is mirrored in other areas of genomics and systems biology, e.g. pseudo-temporal ordering approaches have until recently (Saelens *et al*., 2019) rarely been rigorously tested. But what is also needed are approaches which allow us to assign confidence to inferred networks, or, more specifically, predicted interactions without recourse to a gold-standard (Pratapa *et al*., 2020). This would most likely have to be indirect and based on biological expectations/knowledge related to the scenario under investigation.

One of the potential initial attractions of using a panel of network inference algorithms is that different methods may capture different aspects of the data and in concert may provide a more complete representation of a “true” network of functional relationships among the genes in an organism under a given scenario. While appealing this notion needs to be viewed with caution (Kirk *et al*., 2015b). Combining the most powerful methods by leveraging their individual strengths is possible in principle (Kuncheva, 2004; Madirolas & de Polavieja, 2015; Díez-Pastor *et al*., 2015; Laan *et al*., 2017), but requires us to characterise each method *O*_*i*_ reliably and independently.

## 6 Conclusion

In summary, unless we know the constituency of the model universe, ℳ, or the ensemble of predictors, *O*_*κ*_, we have little way of telling whether we are dealing with a *Madness of crowds* or a *Wisdom of crowds* scenario. However, the present analysis shows that ensemble procedures will be robust as long as the ensembles are carefully constructed. In the context of biological network inference, reduction in false-positives, are the primary cause for their success. Without a robust and transferable way of assessing the strengths and weaknesses of different methods, we cannot (yet) use tools from decision theory that pool these strengths for diverse and heterogeneous ensembles (Madirolas & de Polavieja, 2015; Díez-Pastor *et al*., 2015). Currently the best advice, in light of the analysis carried out here, is to be ruthless in weeding out poorly performing methods for network inference, or models with low weight for multi-model averaging. So there is no need, for example, to include correlation networks, even though they are cheap to calculate: their performance is simply too poor to warrant inclusion in an ensemble. Quality is more important than quantity.

## Software

A Jupyter Notebook containing the Julia code to reproduce all the computational results here, and to explore the effects of the *madness of crowds* in network inference and model averaging is available at https://github.com/MichaelPHStumpf/Madness-Of-Crowds

## A Assessing the performance of network inference methods

Real biological networks are sparse (Lèbre *et al*., 2010); this means that a predictor which scores each candidate edge to be absent can have high performance if the number of false-positives is heavily influencing how we quantity performance of a network inference methods. We therefore focus here on *precision* and *recall* and a derived statistic, the *F*1 statistic (Lèbre *et al*., 2010; Murphy, 2012). We denote the numbers of true and false positive inferences (i.e. scored edges in the context of network inference) by *TP* and *FP*, and the true and false negatives by *TN* and *FN*. Then the *precision, P*, is given by

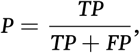

and the *recall* by

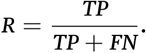

The *F*1 statistic is given by

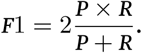

These statistics are confined to the [0, 1] range, with larger values indicating better performance, and the *F*1 statistic in particular becomes maximal at the point where the curves for *P* and *R* intersect.

## B Approximating the Hypergeometric function

The Hypergeometric function (Arfken *et al*., 2013), _2_*F*_1_, appearing in Eqn. (23) can be unwieldy to evaluate for large *k* and *k*_0_ and for sufficiently small of *t* we can Taylor-expand it around *t* = 0 and then obtain (if we restrict the expansion of the Hypergeometric function to third order)

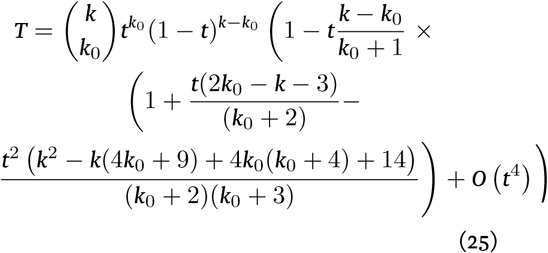

From this we can determine when *T* will be greater than *t* by solving

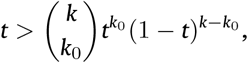

because the term in the brackets in Eqn. (25) is ≤ 1.

## Notes

https://github.com/MichaelPHStumpf/Madness-Of-Crowds

### Competing Interest Statement

The authors have declared no competing interest.

